# Identifying the exposure of taxonomic, functional, and phylogenetic diversity of steppe birds to renewable energy developments

**DOI:** 10.1101/2025.11.07.687200

**Authors:** Pablo Medrano-Vizcaíno, François Mougeot, Beatriz Arroyo, Gerard Bota, David Giralt, Laura Maeso-Pueyo, Carlos A. Martín, Manuel B. Morales, Pedro P. Olea, Juan Traba, Ana Benítez-López

## Abstract

Biodiversity is globally threatened by human impacts including land use transformation and climate change. Global warming has prompted a rapid transition from fossil fuels to cleaner energy sources such as photovoltaic (PV) energy. However, utility-scale PV plants demand vast areas and can lead to conflicts with biodiversity conservation, making strategic planning essential. We evaluate the spatial overlap between conservation priorities and PV infrastructure using steppe birds—a highly threatened bird group occurring in lands potentially suitable for PV plants—as a model. First, we quantified and mapped taxonomic (TD), functional (FD), and phylogenetic diversity (PD) of 26 species across a main stronghold for European steppe birds (mainland Spain and Balearic Islands) to identify prioritization scenarios that best preserved all steppe bird diversity facets (TD, FD, and PD), and steppe birds of conservation concern. Next, we generated a diversity hotspot map based on the combination of TD, FD, and PD and overlapped with existing PV infrastructure to determine: (1) exposure areas (high diversity hotspots with high PV occupancy), and (2) no-go areas (high diversity hotspots with low/null PV occupancy). We found that the prioritization scenario combining TD+FD+PD retained 68.5% of TD, 75.5% of FD, 59.7% of PD, and 62.8% of areas with species of conservation concern, providing a more balanced representation across biodiversity facets than other prioritization scenarios. PV infrastructure currently occurs in 53.1% of high diversity hotspot cells. Overall, exposure areas (7.2% of the study area) were mainly concentrated in Central, Southern, and Northwestern Spain, while no-go areas (19.9% of the study area) were concentrated in Northern, Central, and Central-Western Spain. Our methodology is adaptable to other species, communities and regions, offering a robust framework for balancing development projects with biodiversity conservation. As the transition to renewable energy accelerates, including biodiversity in energy planning is crucial for preventing irreversible ecological losses.

## 1. Introduction

Human activities alter biodiversity, with the most prominent threats being habitat loss caused by land transformation and, more recently, climate change (IPBES 2019; Powers and Jetz 2019). In response, global efforts to reduce carbon emissions have led to a significant and ongoing transition from fossil fuels to cleaner energy sources, such as solar and wind power, which produce lower or no greenhouse gas emissions (Chien et al., 2022). Photovoltaic (PV) solar energy accounted for nearly one-third of the total installed renewable energy capacity in the world in 2022 and its generating capacity has grown over 40% since 2009, with projections of a ten-fold increase by 2040 (Pourasl et al., 2023; Kruitwagen et al., 2021). At the regional level, the European Green Deal aims to reduce greenhouse gas emissions to at least 50% by 2030 compared with 1990 levels and to achieve carbon neutrality by 2050 (European Commission 2019). This plan, updated in 2022, included huge developments in renewable energy, doubling photovoltaic capacity by 2030 (European Commission 2022).

Although renewable energy projects aim to mitigate climate change effects by decarbonization of the energy sector, large-scale solar farms require considerable extents of land, which can lead to conflicts with other important land uses, especially food production and biodiversity conservation (Serrano et al., 2020; Dunnett et al., 2022; Hermoso et al., 2023). Indeed, in regions with high energy demand, such as Europe, India, Japan, and South Korea, PV installations are expected to occupy between 0.5% and 5.2% of the land by 2050 (van de Ven et al., 2021), with concomitant consequences for biodiversity such as habitat loss as a result of rapid habitat transformation. Other negative effects are derived from habitat alterations (removal of vegetation, soil erosion, pollution, and modifications to microclimatic conditions, light patterns, hydrology, and soil properties) or increased animal mortality from various sources, including collisions with PV panels (Dhar et al., 2020; Lambert et al., 2022; Gómez-Catasús et al., 2024). Additionally, the associated infrastructure needed for PV projects, such as roads and power lines, can fragment landscapes, disrupt habitat connectivity, and cause mortality (Guil and Pérez-García 2022; Medrano-Vizcaíno et al., 2023). Indeed, species richness and density have been found to be lower in areas with PV installations compared to adjacent, undisturbed areas (Visser et al., 2019). These ecological consequences can have long-term effects as the lifetime of PV panels is around 20–30 years (Semeraro et al., 2022), and they might be renewed after this time.

In turn, some species may benefit from solar farms when wildlife needs are considered (e.g. plants in hydrologically deprived environments, Liu et al., 2019; birds in arable farmlands, Copping et al., 2024). Nonetheless, the scientific evidence on PV impacts on biodiversity, whether negative or positive, is still sparse, focused on local-scale (e.g. landscape and microhabitat) impacts, and geographically and taxonomically biased towards the USA and UK, and arthropods and plant species (Lafitte et al., 2023, Gómez-Catasús et al., 2024). Studies on vertebrate species with large-scale ecological requirements and studies linked to habitats with topographic and climatic features targeted by PV developers are missing.

Assessing the impacts of renewable energy projects on biodiversity is crucial to ensure that sustainability and conservation goals are met (Gómez-Catasús et al., 2024). A first step to avoid PV impacts is the implementation of proactive land-use planning at regional or national scales, using ecological attributes for providing recommendations for spatial siting of renewable energy projects in areas that do not conflict with biodiversity (Hermoso et al., 2023). A wealth of research has focused on how PV systems intersect with Important Conservation Areas (ICAs), such as protected areas, key biodiversity areas, and wilderness areas (Santangeli et al., 2016; Sonter et al., 2020; Dunnett et al., 2022; Niebuhr and Sant 2022). Yet, these assessments fail to recognize that impacts on species and functional groups are uneven (Pérez-García et al., 2022). For instance, highly mobile species such as birds can spend long periods outside ICAs (Boakes et al., 2019) and suffer anthropogenic mortality due to collision with infrastructures. In the United States, 9.9 birds/MW are estimated to die each year due to collisions with solar facilities (Walston et al., 2016). Additionally, as grasslands and agricultural lands tend to be less protected than forests, species linked to these habitats spend much of their life cycle outside ICAs, hence being disproportionately exposed to PV developments that may not have to comply with regulations linked to ICAs (Serrano et al., 2020). This issue deserves particular attention, as even some steppe bird species (e.g., *Pterocles orientalis*) have been reported to become locally extinct due to poor PV plant planning (Bolonio et al., 2024).

Steppe birds are declining worldwide and are the most threatened group of birds in Europe (Burfield et al., 2023). These birds may be particularly vulnerable to PV developments as they inhabit low-intensity flat agricultural areas, where many PV projects are currently being developed (Serrano et al., 2020). This scenario is exacerbated in Spain, a territory that contains a large proportion of the European steppe bird populations and that constitutes one of their main strongholds (Burfield et al., 2023, Medrano-Vizcaíno et al. 2025). Here, remarkable declines have been observed in recent years (Traba and Morales, 2019; Traba and Pérez-Granados 2022), reflected in notable changes in the threat status at the country and continental level for several species (Medrano-Vizcaíno et al. 2025; Gómez-Catasús et al., 2025). Indeed, the comparison of the Spanish Red Lists (SRL) of 2004 and 2021 (Madroño et al., 2004; SEO/BirdLife 2021) shows that 13 (50%) out of 26 steppe bird species have been upgraded to a higher conservation status. The threat status of steppe bird species could be further worsened by the Spanish National Integrated Energy and Climate Plan (PNIEC) for 2023–2030 (MITECO, 2024), which plans an increase of approximately 65,000 MW of solar photovoltaic energy production by 2030 in comparison with 2020. While there exist non-binding tools to provide information to developers that facilitate biodiversity-friendly renewable developments (Environmental zoning of renewable energies: Wind and Photovoltaic, MITECO, 2020), these are focused on the network of Important Bird Areas, the Natura 2000 network, and other areas of environmental significance, thus neglecting low-productivity agricultural lands and grasslands, which usually host steppe bird populations (Medrano-Vizcaíno et al. 2025). Therefore, a novel and refined prioritization tool that informs photovoltaic siting is key to reducing impacts on threatened steppe bird communities.

To gain a deeper understanding of the ecological processes that shape biodiversity and to refine conservation priorities, biodiversity should not be understood solely in terms of taxonomic diversity (TD) (Pollock et al., 2017). A thorough assessment must include other facets, such as evolutionary history (phylogenetic diversity, PD) and functional diversity (FD). PD provides insights into the evolutionary uniqueness of species within a community (Lum et al., 2022), while FD focuses on the variety of functional traits in communities, which can be useful in assessing ecosystems’ resilience, stability and functioning (Sol et al., 2020). An integrated analysis of TD, PD, and FD can help identify conservation-valuable areas and potential conflicts with human activities from a more complete picture of biodiversity (Smiley et al., 2020). This approach can help pinpoint areas at high risk for biodiversity loss and guide decisions about where stricter protections are needed, being especially important for highly threatened communities.

We analyse steppe birds as a case study in the context of the current worldwide expansion of photovoltaic energy, by addressing two key questions: 1) Does a multifaceted prioritization approach (TD+FD+PD) outperform single-facet strategies in retaining high biodiversity areas and areas with species of conservation concern?, and 2) To what extent do PV infrastructures spatially overlap with steppe bird diversity hotspots (TD, FD, PD) in Spain?

For this, we leveraged the most recent data on the occurrence of 26 steppe bird species in Spain (III Spanish Atlas of Breeding Birds; SEO/BirdLife 2022; Medrano-Vizcaíno et al. 2025) to identify high diversity areas based on multiple biodiversity facets (taxonomic – TD, functional - FD, and phylogenetic diversity - PD), thus ensuring the protection of individual species, but also of evolutionary distinctiveness and the diversity of the functional traits that support important ecosystem functions. We quantified the coverage of each facet that is retained when prioritized individually compared to when they are combined, thereby highlighting potential trade-offs and complementarities among biodiversity dimensions. We also evaluated the effectiveness of each prioritization scenario (single-facet and combined) in retaining areas with steppe birds of conservation concern. Finally, we overlapped the best prioritization scenario with a newly generated spatially explicit map of installed photovoltaic infrastructures across Spain to identify 1) exposure areas, where high diversity areas coincide with high PV occupancy, and 2) no-go areas, that is, high diversity areas with low or null PV occupancy, and where future PV implantation should be avoided or minimized. Our results provide a framework that derives generalizable conclusions and recommendations and may be transferred to other taxa and regions.

## 2. Methods

### 2.1 Study area and species occurrence

We performed our analyses across Peninsular Spain and the Balearic Islands, a territory that contains a great proportion of European steppe birds (Burfield et al., 2023). In particular, Iberia harbours steppe bird species distributed over different regions in the Palearctic, like the Mediterranean basin, Atlantic Europe or central Asia, which confers high evolutionary and biogeographical singularity to the Iberian steppe avifauna (Santos & Suárez, 2005). As spatial units, we used Universal Transverse Mercator (UTM) grid cells of 10 × 10 km, which totaled 3,838 cells that contained at least 5 species, which we considered the minimum number for meaningful FD analyses (see below).

We used occurrence data for 26 species (Appendix 1) that have also been included in previous steppe-bird conservation research in Spain (Medrano-Vizcaíno et al., 2025, Traba et al., 2007). These species are typical of Iberian natural steppes and agricultural pseudo-steppes (Suárez et al., 1997; Traba et al, 2007), are essentially ground-nesting, occur in treeless and mainly flat areas, and have their main European populations in the Iberian Peninsula (Medrano-Vizcaíno et al., 2025). As for steppe habitats, we refer to natural and semi-natural open habitats, including natural steppes, dry cereal farmlands, grasslands, and shrublands (Sainz Ollero, 2013).

Data was compiled from the III Spanish Atlas of Breeding Birds (period 2014-2018) (SEO/BirdLife 2022) and eBird (records during the breeding period: February-August 2019-2023), therefore covering a decade (see more details in Medrano-Vizcaíno et al., 2025).

### 2.2 PV energy data

We searched and gathered all available geospatial PV databases at global, national, and regional levels which consisted of polygons or points, or both. We completed this information with the SatlasPreTrain Geospatial Data (Bastani et al., 2023), which consists of a high-accuracy AI-generated output of polygons classified as solar farms at a global scale. Additionally, we identified areas where spatially-explicit data on PVs were missing based on the Spanish PRETOR database (MITECO, n.d.), which contains information on the installed PV power per municipality. Missing PVs were georeferenced with the aid of high-resolution PNOA - Plan Nacional de Ortofotografía Aérea - images (Instituto Geográfico Nacional, n.d.), adding points separated by ca. 250 m and ensuring that PV plant areas were homogeneously covered. Finally, we merged all the different point databases and applied a 250 m buffer around PV points so that the whole area of the solar PV plant was covered. Using these polygons, we constructed a spatial layer depicting the proportion of cell area covered by PV plants.

### 2.3 Biodiversity facets analysis

Taxonomic Diversity (TD) was calculated as the number of species present in each cell. Functional Diversity (FD) was calculated using the FRic (functional richness) index, which quantifies the amount of functional space occupied by the species of a given community (Villéger et al., 2008). This index is more accurate than the dendrogram-based index and is comparable with species richness and the Faith’s PD index for phylogenetic diversity, as all three reflect the extent of diversity based on richness, whether taxonomic, functional, or phylogenetic (Maire et al., 2015; Tucker et al., 2017). To this end, for each species, we gathered data on 32 traits related to morphometry, body mass, life history, habitat, diet, population density, and migratory behaviour from different sources such as databases (e.g., Tobias et al., 2022, Cooke et al., 2019, Santini et al., 2023), books (e.g., Madroño et al., 2004, Snow and Perris 2008), scientific papers (e.g., Tsuboi et al., 2018; Pérez-Granados et al., 2013), as well as our own field data (see Appendix 1 for the complete list of species and traits). Nevertheless, we dealt with missing data for certain traits: birth or hatching weight (10 missing values), brain size (8 missing values), habitat breadth (2 missing values), degree of development (1 missing value), and population density (3 missing values). To avoid removing traits or species with incomplete data, we applied imputation methods, which rely on the fact that traits are usually correlated, to complete missing values (Johnson et al., 2021). Imputation was performed using the R package function *missforest*, an approach that performs better than other methods such as k-nearest neighbours imputation or multivariate imputation (Stekhoven 2013; Tucker et al., 2017).

Later, we built a species × trait matrix (i.e., 26 rows x 32 columns), and a community × species matrix (i.e., 3,838 rows x 26 columns). This latter matrix represents the presence/absence of each species in each grid cell. We used a minimum threshold of 5 species per cell because the number of species in each one must be higher than the number of dimensions used to define the multidimensional functional space based on PCoA axes (we selected four dimensions, which accounted for 88% of the total variance based on the relative eigenvalues; Table S1). As including additional dimensions would have resulted in a significant loss of data (more grid cells excluded from analyses), the selection of four dimensions ensured a meaningful representation of the functional space while retaining as many cells as possible, thus minimizing data loss.

Given that the FRic index is highly influenced by TD (i.e., a higher number of species can lead to a broader range of traits, depending on species trait variation) (Plass-Johnson et al., 2016), we calculated the Standardized effect size of FRic (SESFRic) to remove the effect of species richness. SESFRic was calculated as observed FRic - mean of expected FRic / SD of expected FRic, where “observed FRic” represents the observed values of FRic, while “mean of expected FRic” and “SD of expected FRic” represent values obtained from null models (Mason et al., 2012). To this end, we executed 1000 permutations of the community matrix using the “independent swap” null model, which randomizes the data matrix and maintains the original species richness (Aros-Mualin et al., 2021). We used the function *dbFD* from the R package *FD* (Laliberté et al., 2014) to calculate FRic, and the function *randomizeMatrix* from the R package *picante* (Kembel et al., 2010) to generate the null models.

To calculate phylogenetic diversity (PD), we used the bird phylogeny of Jetz et al. (2012) and applied the Faith’s PD index, which represents the sum of phylogenetic branch lengths connecting species within a community (Faith 1992). As for FRic, this index is highly sensitive to TD, therefore we used the function *ses.pd* from the R package *picante* to calculate the effect size of PD (SESPD) based on 1000 permutations that randomize species labels while preserving the underlying phylogenetic structure. The use of standardized effect sizes can minimize correlations among biodiversity facets (see Table S2).

To allow comparability across analyses, all three biodiversity facet values: TD, SESFD (hereafter FD), and SESPD (hereafter PD) were scaled between 0 and 1 by subtracting the minimum and dividing by the difference between the maximum and the minimum. As an additional analysis we assessed the contribution of each species to the overall FD, and PD. For this, we iteratively removed each species and recalculated the resulting FD and PD. The contribution of each species was computed as the difference between the total FD/PD using all species and the FD/PD obtained without that species.

### 2.4 Spatial prioritization scenarios

We used the spatial prioritization software Zonation 5 (Moilanen et al. 2022) to generate four different prioritization scenarios, three of them representing single biodiversity facets: TD, FD, and PD, and another one combining all three facets: TD+FD+PD (where TD, FD, and PD had the same importance by assigning an equal weight of 1). Zonation 5 produces a hierarchical prioritization of the study region sites (cells) based on their biological value, accounting for complementarity by considering the representation level of biodiversity features (i.e., biodiversity facets in our study) (Moilanen et al., 2022). We selected the Core-Area Zonation 2 (CAZ2) marginal loss rule, which enhances coverage for underrepresented features at the cost of a slight reduction in overall average coverage. This rule has shown better performance than other marginal loss rules, such as ABF (Additive-Benefit Function) and CAZMAX (Core-Area Zonation): while ABF prioritizes higher average coverage at the expense of underrepresented features, CAZMAX results in lower overall average coverage (Moilanen et al., 2022).

To assess the effectiveness of each scenario in preserving TD, FD, and PD, we overlapped the top 30% of TD, FD, and PD cells (separately) over the top 30% of each prioritization scenario and calculated the retention of the three biodiversity facets under each prioritization scenario. This was useful to quantify to what extent taxonomic, functional, and phylogenetic diversity are preserved under each of the four scenarios.

We also analysed the coverage of these scenarios over areas inhabited by steppe bird species of conservation concern (threatened species areas, hereafter). To this end, we used the occurrence data of the 26 steppe bird species to derive a threatened species index map based on the combination of three threat status categories at European and Spanish level: the European Population Status (EPS) (BirdLife International, 2004; Burfield et al., 2023), the Species of European Conservation Concern (SPEC) (BirdLife International, 2004; Burfield et al., 2023), and the Spanish Catalogue of Threatened Species category (hereafter SCTS), which is the official and legal instrument for the protection of wild species in Spain (MITECO, 2023). Following Medrano-Vizcaíno et al. (2025), we used a numeric scoring scale that ranged from 0 to 10, where 10 represents the highest concern category (Table S3). The individual scores were summed across species per grid cell and standardized from 0 to 1. Finally, the standardized scores were summed to generate the threatened species index per cell. Next, we overlapped the top 30% of each prioritization scenario over the top 30% of the threatened species index (i.e., threatened species areas) (Figure S1) to calculate the coverage of each prioritization scenario over regions hosting a high number of threatened species. The best prioritization scenario was selected based on the highest average retention and more balanced values across facets.

We focused on the 10 × 10 km grid cells with the top 30% values of each biodiversity facet, aligning with the targets set by the Kunming-Montreal Global Biodiversity Framework (2022) and the EU Biodiversity Strategy (2020), both of which advocate conserving at least 30% of global and European land areas by 2030.Since all grid cells represent equal areas, the top 30% of values corresponds approximately to 30% of the study area, providing a spatially consistent way to identify priority areas for biodiversity conservation.

### 2.5 Steppe birds’ exposure to photovoltaic facilities

To assess the current and potential exposure of steppe birds to photovoltaic (PV) infrastructures, we overlaid the diversity hotspot map generated by Zonation 5 (i.e., TD+FD+PD scenario) with PV occupancy (the proportion of land within each grid cell currently covered by PV infrastructure). Both diversity hotspots and PV occupancy values were divided into tertiles (low, medium, high). PV occupancy was standardized to a 0–1 scale (min-max scaling) to align with the range of the diversity hotspots map. Original PV occupancy ranged from <0.01% to 13.14%, with high occupancy defined as 2.85–13.14%, medium as 0.32–2.77%, and low as <0.01–0.32%.

The combination of the PV occupancy and diversity hotspots maps resulted in nine joint categories (i.e., high diversity hotspots - high PV occupancy, high diversity hotspots - medium PV occupancy, etc.) that were represented using bivariate maps using the *biscale* package in R (Prener et al., 2022). We identified as *exposure areas* those cells with high values (upper tertile) of both diversity hotspots and PV occupancy maps. On the other hand, cells with high diversity hotspots values and low PV occupancy were identified as *no-go areas* where PV developments should be avoided or minimized. These areas represent locations with minimal or no current PV infrastructure but where future developments could have significant impacts on steppe birds.

We present our results both at the national level and at the level of administrative regions because, in Spain, the latter have the legal competences on environmental policy and PV approval when plants have an installed capacity of < 50 MW (IEA, 2021).

All analyses were conducted with R Statistical Software (v4.2.2) and RStudio (v2023.12.1.402), ArcGIS (v.10.4.1) and Zonation 5 (v.2.1).

## 3. Results

We found marked differences in spatial patterns among biodiversity facets (Figure 1). Notably, a high diversity of any facet did not imply high diversity for other facets, with Pearson correlation coefficients ranging between - 0.1 and 0.5 (Table S2). For example, northern Spain, and some regions in western Spain showed a high TD only, while regions in southern Spain showed particular areas with an increased PD only. Likewise, some regions in Central Spain showed areas with high FD only. Also, insular areas appeared more important under the PD scenario. The contribution of individual species to both FD and PD varied considerably (Figures S2 and S3). The Great Bustard *Otis tarda*, the Zitting Cisticola *Cisticola juncidis*, and the Eurasian Thick-knee *Burhinus oedicnemus* showed the highest contributions to FD (i.e., their removal resulted in the largest FD reduction). Regarding PD, the species with the highest contributions were the Short-eared Owl *Asio flammeus*, the Eurasian Thick-knee, and the Lesser Kestrel *Falco naumanni*.

**Figure 1.**
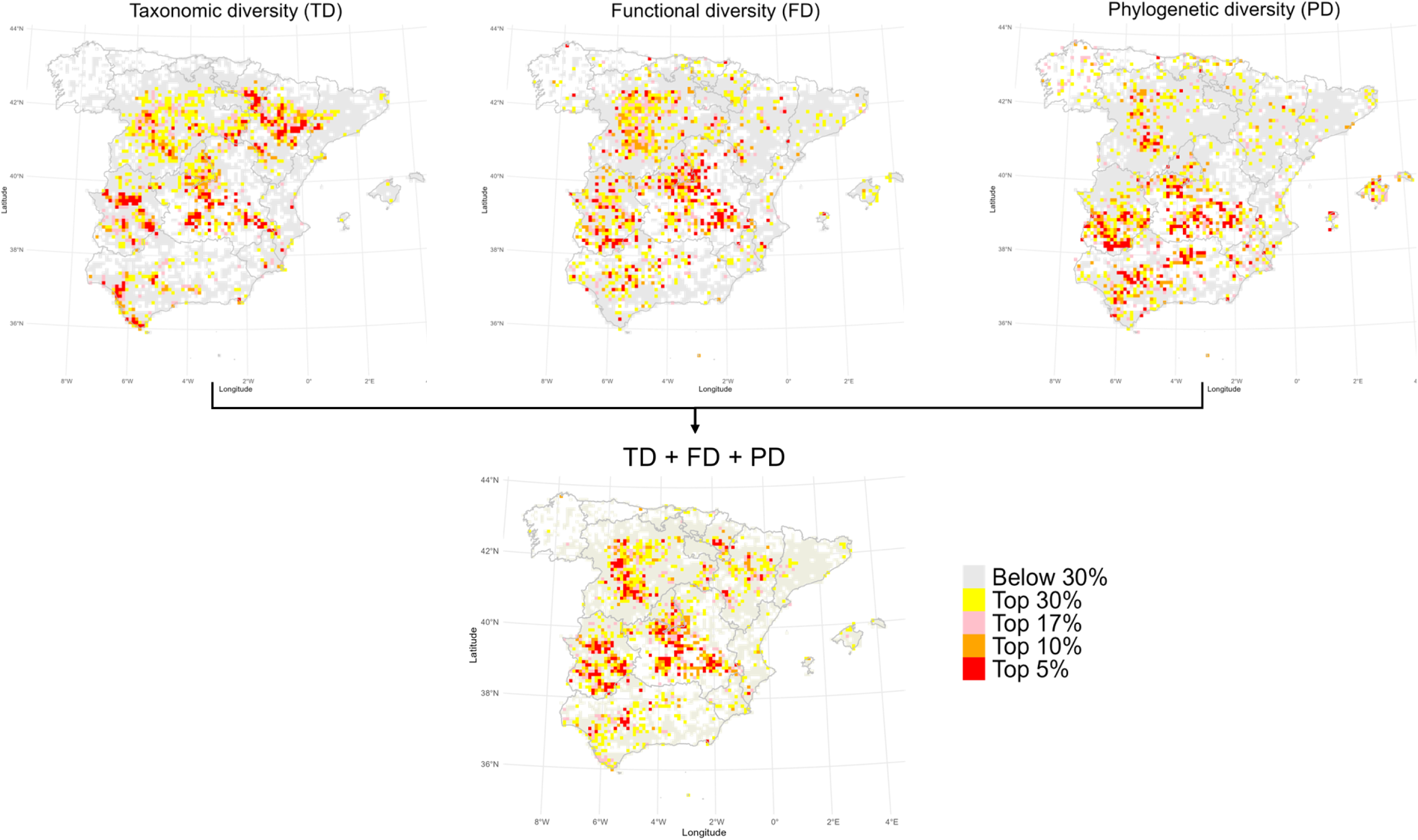
Prioritization maps under taxonomic, functional, phylogenetic diversity approaches separately, and under the scenario combining all three diversity facets (Top 30%, 17%, 10%, and 5%) in peninsular Spain and Balearic Islands. We show the top 30%, 17%, 10%, and 5% priority areas to facilitate comparison among scenarios and to align with key international conservation targets. Specifically, the 30% and 17% thresholds reflect the goals of the Kunming-Montreal Global Biodiversity Framework (2022) and the EU Biodiversity Strategy (2020), which call for the protection of at least 30% of terrestrial areas by 2030, and the Aichi Target 11 from the Convention on Biological Diversity, which aimed to conserve at least 17% of terrestrial land by 2020 (CBD, 2010).

According to Zonation 5, the most inclusive conservation scenario was TD+FD+PD, as it covered areas that remained unprioritized in the other scenarios. This scenario included: areas in central and western Spain that were not highly prioritized under the PD scenario; areas in northeastern Spain that were not prioritized under the FD and PD scenarios; and some areas in central Spain that were not prioritized under the TD and PD scenarios. Interestingly, regions along the northern and northwestern borders of the country were not prioritized under any scenario (Figure 1).

We found that the TD+FD+PD scenario retained the highest overall proportion of biodiversity facets and areas inhabited by threatened species (Figure 2). This scenario achieved an overall retention of 66.6%, including 68.5% of TD, 75.5% of FD, 59.7% of PD, and 62.8% of threatened species areas. It was followed by the TD scenario, with an overall retention of 64.3%, although inflated by its evident 100% coverage of TD, and showing poor representation of FD (50.1%) and PD (33.9%), despite high coverage of areas with threatened species (73.3%). The FD scenario performed worse, retaining 50.1% of TD, 54.2% of PD, and 49.4% of threatened species areas. The PD scenario showed even lower and more unbalanced retention percentages, with 33.9% of TD, 54.2% of FD, and 37.1% of threatened species areas. Overall, the TD+FD+PD scenario also achieved the highest average retention and more balanced values across facets when compared with the two-facet scenarios (Table S4).

**Figure 2.**
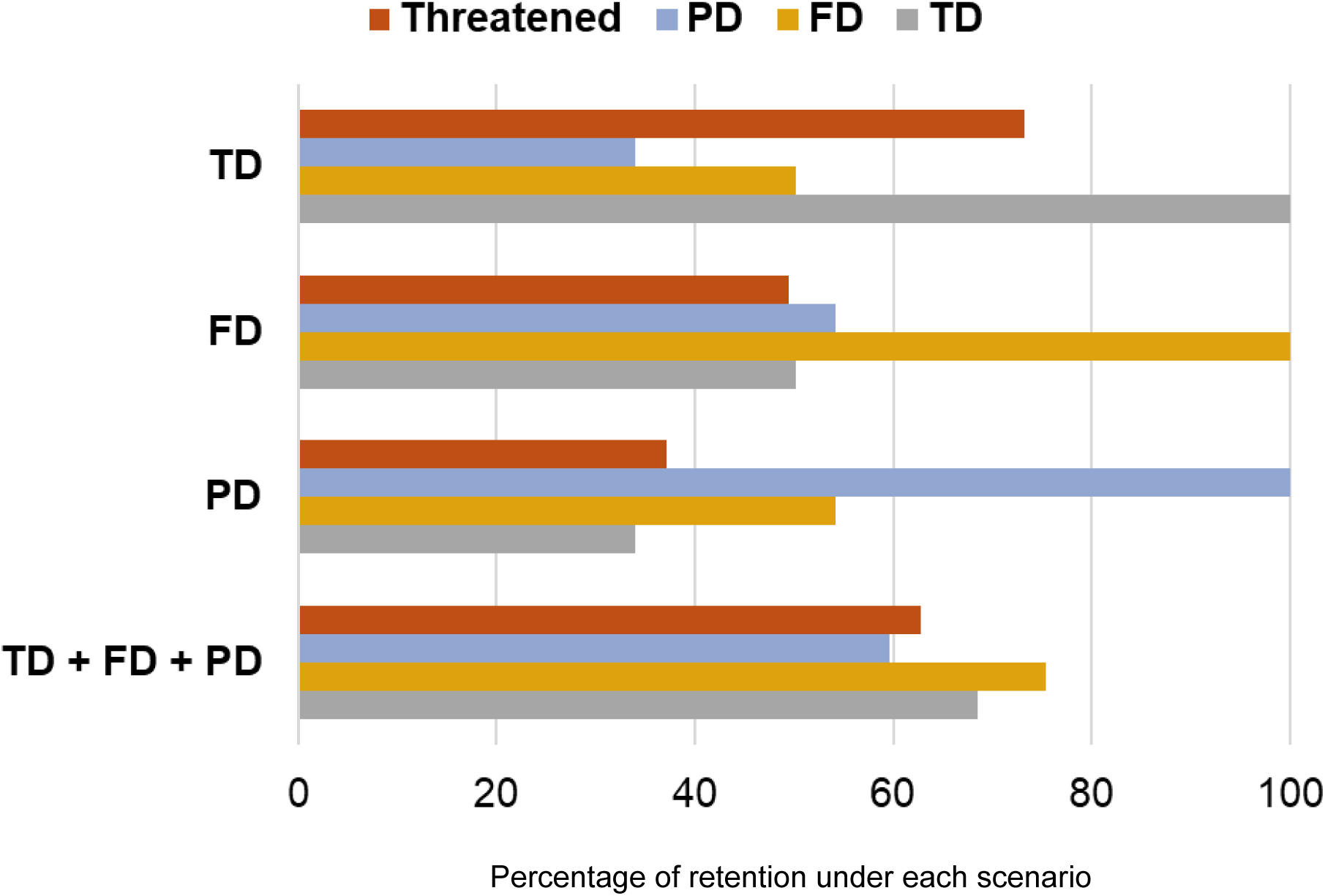
Percentage of each biodiversity facet and the *threatened species index* retained under the four prioritization scenarios. The y-axis represents each prioritization scenario, and the x-axis is the percentage of each biodiversity facet, and the *threatened species index* retained under each prioritization scenario.

### 3.1 Steppe birds’ exposure to solar photovoltaic facilities, and no-go areas

We found that 40.5% of all cells and 53.1% of high diversity hotspots cells host PV facilities. The intersection between the diversity hotspots and PV occupancy maps revealed distinct patterns of exposure of steppe bird diversity to solar energy development (Figure 3). We detected 276 exposure cells having both high diversity hotspots and high PV occupancy (7.2% of our study area). Exposure cells were mainly concentrated in central, southern, and northwestern Spain. Here, steppe bird communities are already under pressure from existing PV development, highlighting potential biodiversity-energy conflicts that require conservation attention.

**Figure 3.**
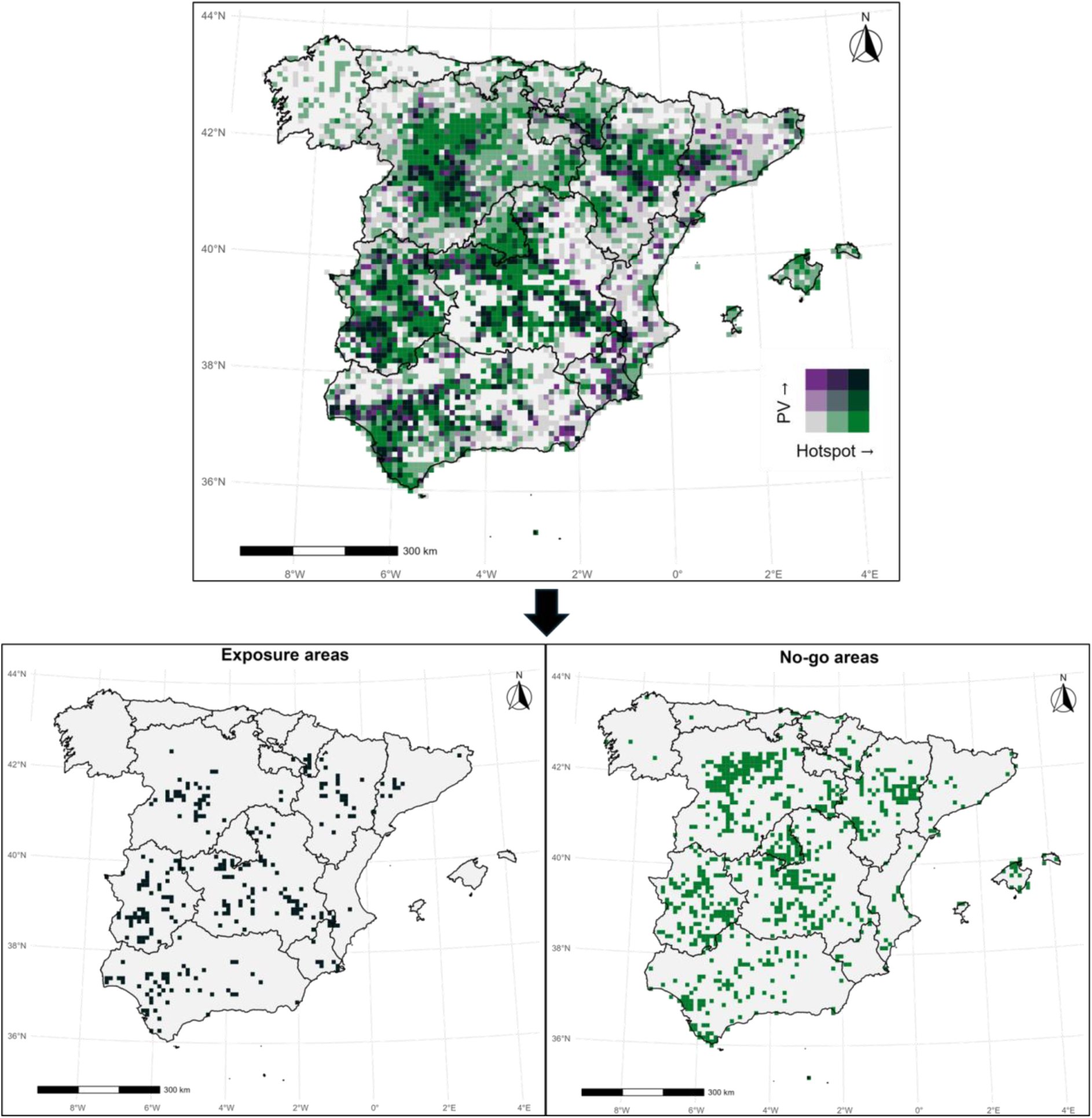
Proposed approach to identify priority areas for the conservation of steppe birds in peninsular Spain and Balearic Islands. The top figure shows a bivariate map combining low, medium, and high hotspots values (TD, FD, and PD combined) and PV (photovoltaic) occupancy. The bottom maps show exposure areas and no-go areas extracted from the bivariate map.

In addition, we identified 765 cells having high diversity hotspots but low PV occupancy (19.9% of our study area), being broadly distributed across Spain; although they showed higher concentration in northern, central, and central-western Spain. These no-go areas currently experience little or no PV development but represent locations where future infrastructures could pose risks to steppe bird communities if not carefully managed.

## 4. Discussion

The rapid global expansion of renewable energy infrastructure is transforming landscapes at an unprecedented pace, often at the expense of biodiversity. The conversion of natural habitats for renewable energy projects poses a critical threat to ecosystems worldwide, exacerbating the ongoing biodiversity crisis (IPBES, 2019). Reconciling biodiversity conservation and climate change mitigation goals is a societal and political challenge. We address this global issue by examining spatial conflicts between PV development and steppe bird conservation in Spain, one of the main strongholds for these species. Our multidimensional analysis, integrating taxonomic, functional, and phylogenetic diversity, revealed numerous regions where high biodiversity value coincided with substantial PV infrastructure, highlighting areas at risk from current and future PV developments. These findings underscore that relying on a single biodiversity facet can overlook critical conservation priorities. By adopting a multidimensional approach, our study provides a robust framework for identifying potential conflicts between renewable energy projects and biodiversity conservation, offering a scalable strategy applicable to threatened ecosystems worldwide. Additionally, the proposed approach can be extended to other infrastructures (e.g. wind farms) and species groups.

### 4.1 Spatial patterns of steppe-bird biodiversity facets

Previous studies have emphasized the role of ecological and evolutionary processes in shaping biodiversity patterns across geographic regions (Monnet et al., 2014). This aligns with our results showing that, although central and western regions within Iberia are taxonomically, functionally, and phylogenetically highly diverse in terms of steppe birds, other regions such as parts of the northeastern interior emerged as important mainly for taxonomic diversity. In contrast, several areas in Southern Spain stand out primarily for their high phylogenetic diversity. These patterns suggest that some areas, although rich in species, support communities composed of species that are functionally similar and evolutionarily close. This is indicative of low ecological differentiation and limited evolutionary distinctiveness, which may reduce the resilience of communities to environmental change (Cadotte et al. 2012). For example, areas with high TD but low FD and PD might be characterized by a prevalence of species-rich taxonomic groups (e.g., certain families of passerines) that, while numerous, share similar functional traits and a relatively recent common evolutionary history. This can result in higher functional redundancy (where many species perform similar ecological roles) and shorter overall phylogenetic branch lengths, meaning the loss of any single species in such a community might result in a comparatively small reduction in functional volume or phylogenetic distinctiveness. On the other hand, areas concentrating high values of all three biodiversity facets, represent critical conservation priorities, as they host not only many species but also deeper evolutionary lineages and a broader range of ecological strategies, from large-sized ground-dwelling herbivores (Great and Little bustards) to small-sized invertivores (Dupont’s lark or Wheatears) and medium-sized raptors (harriers).

Recognizing these multidimensional patterns is essential for anticipating how different human pressures may impact biodiversity in varied ways. For example, intensive agriculture has been shown to reduce functional diversity in farmland birds (Guerrero et al., 2024), a concern for steppe birds that largely rely on cereal steppes. Hunting pressure, habitat destruction, and land-use changes have been associated with declines in taxonomic, functional, and phylogenetic diversity, respectively (Romero-Muñoz et al., 2021). Similarly, climate change is expected to negatively affect functional diversity (Penjor et al., 2022), while urbanization poses risks particularly to phylogenetic diversity (La Sorte et al., 2018). Recently, Gómez-Catasús et al. (2025) identified that threatened European steppe birds are mostly large-sized, long-lived, ground-dwelling and ground-foraging species, providing insights on how agriculture intensification and human-induced mortality are eroding non-random sections of their functional space.

While recognizing the different patterns of biodiversity can be key to understanding the underlying processes that act on maintaining ecosystems (Rosauer et al., 2009), a multifaceted approach emerged as more effective for steppe bird conservation prioritization.

### 4.2 Spatial Prioritization

Protecting multiple facets of biodiversity presents significant challenges, as it often involves unavoidable trade-offs (Wilson & Law, 2016). No prioritization scenario was able to fully preserve all three biodiversity facets. However, our results show that integrating taxonomic, functional, and phylogenetic diversity maximizes the overall coverage of biodiversity facets and areas with high threatened species index. This scenario (TD+FD+PD) retained almost 60% of PD, 75% of FD, nearly 70% of TD, and more than 60% of regions with high threatened species, capturing the complex interactions between threatened species, their ecological roles, and their evolutionary history, key factors for conservation planning (Brum et al. 2017, Pollock et al. 2017).

Our results demonstrate that prioritization scenarios based on a single biodiversity facet were not effective at preserving other facets. However traditional conservation strategies and legislation have focused predominantly on TD, neglecting functionally and evolutionarily diverse regions (Cadotte & Tucker, 2018). For example, the TD scenario showed the worst performance among all scenarios, covering only one-third of PD and half of FD. This highlights a critical issue: a TD-centric approach can overlook key areas that, while perhaps not maximizing species numbers, are crucial for other biodiversity dimensions. For instance, such an approach might undervalue regions where the presence of species with unique functional traits (e.g., the Great Bustard, Zitting Cisticola, and Eurasian Thick-knee) is vital for maintaining functionally diverse ecosystems. Similarly, areas harboring species that represent particularly distinct evolutionary lineages (e.g., the Short-eared Owl, Eurasian Thick-knee, and Lesser Kestrel, as evidenced in the community’s phylogenetic tree; Figure S4) could be missed, leading to a significant loss of phylogenetic diversity. These examples illustrate a broader principle applicable to many ecosystems: conserving the full spectrum of biodiversity requires looking beyond simple species counts.

Conservation programs are often required to balance competing demands, such as habitat restoration, species monitoring, and the mitigation of human impacts, all while adhering to financial constraints, and insufficient political will. In the face of these challenges, a key consideration for conservationists is how to effectively allocate limited resources to achieve the greatest possible benefits. Our prioritization approach, while emphasizing areas with high biodiversity across the three facets, also aimed to improve coverage of underrepresented biodiversity facets across grid cells by using the CAZ2 marginal loss rule. This differs from more restrictive prioritization approaches in Zonation 5 software (e.g., ABF, and CAZMAX), which have a lower distribution coverage of underrepresented features (in this case biodiversity facets) (Molainen et al., 2022). These latter prioritization approaches would have, for instance, excluded areas with high FD and PD but low TD. Conserving these areas, where species richness may be low but harbor species with unique evolutionary lineages that play key ecological roles, is crucial, as their loss could cause disproportionate ecosystem disruptions (Rosauer et al., 2009). These findings highlight the necessity of a multidimensional conservation approach to ensure the long-term sustainability of species assemblages and ecosystem functions. While identifying priority areas is a crucial step, translating this knowledge into effective spatial planning is equally important, especially in the context of rapid PV expansion.

### 4.3 Conflicts between steppe-bird conservation and photovoltaic (PV) Infrastructures

Strategic planning is urgently needed to balance renewable energy expansion and biodiversity conservation. In Spain, the rapid growth of PV energy is increasingly threatening steppe bird habitats, posing a major risk to already declining steppe bird populations (MITECO, 2024; Serrano et al., 2020).

Our analysis provides a clear framework for identifying areas where PV developments could threaten steppe-bird biodiversity, allowing for informed decision-making. The representation of different levels of diversity hotspots and PV occupancy to identify exposure and no-go areas is important because the effects of PV developments are not uniform across all areas. For example, while the area occupied by PV plants in some regions may seem small, the localized effects on biodiversity can be severe. Unless conservation planning is effectively implemented there is a high risk that high diversity areas will be exposed to meet the 76000 MW target by 2030 (57000 MW of utility-scale photovoltaics and 19000 MW of self-consumption, MITECO, 2024).

Steppe birds are already under considerable pressure due to agricultural intensification, abandonment of extensive grazing, and human-induced mortality due to agricultural practices and collisions with powerlines, factors that have led their populations to notably decline worldwide (Traba & Morales 2019; Traba & Pérez-Granados, 2022; Gómez-Catasús et al. 2025). An uninformed and disproportional PV infrastructure expansion can be an additive highly threatening factor, especially in Central, Central-Western, Northern interior, and Southern Spain, where we identified most current exposure and no-go cells. Indeed, some regions in Southern and Central-Western Spain have the highest installed capacity and generated PV power in Spain (Red Eléctrica de España, 2025). Without targeted conservation efforts, large-scale PV developments in these areas could result in substantial habitat loss, increased mortality due to collisions with PV panels and distribution lines, and disruptions to ecosystem functions (Dhar et al., 2020; Lambert et al., 2022; Moscatelli et al., 2022).

We consider that periodical conservation assessments and targeted habitat restoration initiatives, particularly in identified exposure areas (where PV infrastructure is already installed, and where potential impacts on birds may be ongoing), could be beneficial for wildlife. Concurrently, policies limiting PV developments in both exposure and designated no-go areas are crucial for safeguarding steppe bird populations.

However, achieving a balance between energy development and biodiversity conservation is necessary. This involves not only careful planning in potentially conflicting areas but also identifying and incentivizing PV development in locations with lower ecological value. An ideal approach could integrate our prioritization findings with “ecovoltaic” facilities and the strategic siting of PV plants on degraded lands (e.g., former quarries, mines, landfills) or within the urban areas (e.g., rooftops of industrial or commercial buildings), thereby minimizing new incursions into sensitive habitats. Ecovoltaic facilities are designed based on ecologically informed strategies that balance energy generation with the preservation and enhancement of ecosystem services (Sturchio and Knapp, 2023), focusing on principles like site selection, park layout, ecosystem design, and adaptive ecosystem management (Tölgyesi et al., 2023). Although no studies have analysed the effectiveness of ecovoltaics on any species, our results offer valuable insights for tailoring such approaches to enhance their potential benefits for steppe bird conservation, contingent on evaluation of their effectiveness. For example, unprotected areas that hold steppe bird populations, but where there is prospective development of solar plants, offer a unique opportunity to implement ecovoltaics in the context of a pilot study that assesses their effects on the behaviour, physiology and/or demography of steppe birds.

### 4.4 Limitations and future research

Despite the comprehensive nature of our study, there are several limitations that warrant attention. First, the responses of species to PV infrastructure can vary depending on local ecological conditions, species behavior, and the magnitude of infrastructure. There is a major knowledge gap in understanding how different species respond to the installation of PV plants in terms of direct impacts (e.g., habitat loss and collisions) and indirect effects (e.g., changes in food availability and provision of breeding habitats). Understanding the interplay of PV plants with variables such as space use, reproductive success, physiological levels or type of flight can provide additional and relevant clues for conservation.

Another important aspect is the evaluation of the potential impacts of emerging energy sources and associated infrastructures (Vermeylen, 2010). Such efforts are essential to understand the cumulative effects of renewable energy development on biodiversity. Importantly, these impacts may not occur in isolation but rather interact synergistically with other well documented threats to steppe birds, such as agricultural intensification, habitat loss, power line collisions, hunting, and climate change (Serrano et al., 2020; Gómez-Catasús et al. 2025). Assessing these combined pressures is critical for designing effective, evidence-based conservation strategies that can mitigate the compounding effects on already vulnerable species and ecosystems.

To our knowledge, our database of PV infrastructure is the most accurate to-date in Spain. We are confident that it captures most PV plants > 5 MW and many more with 1 MW installed power. Ideally, this information should be provided and updated regularly by regional governments, which is essential in the case of Spain because the regional governments hold the responsibility to approve PV developments, as well as the responsibility of preserving vulnerable species. In any case, the spatial analysis of our national-scale PV database, together with data on a threatened group such as steppe birds, highlights opportunities to minimize potential biodiversity impacts and for strategic energy planning.

Our approach can be directly integrated into Spain’s policy and regulatory frameworks for PV development. Currently, PV sites are selected based on factors favorable to developers (e.g. grid connections, solar potential, and land availability). As environmental impact assessments are mandatory (BOE, 2023), our prioritization maps can guide regional authorities during early site evaluation. This tool complements and refines MITECO’s environmental zoning system, aligning with the impact mitigation hierarchy (Phalan et al., 2018). At broader scales, it can inform regional and national energy policies such as REPowerEU, the Fit for 55 package (European Commission 2022), and the PNIEC 2023–2030 (MITECO 2024), which is the national strategic plan that sets out the roadmap for climate and energy policy until 2030. More immediately, our maps provide a valuable resource for designating Renewable Acceleration Areas, required by early 2026 (European Commission Parliament, 2022).

Nevertheless, we acknowledge that the spatial scale used and the quality and phenology of the data (Atlas and eBird data, which do not arise from standardized sampling in the whole territory and may include information gaps), may be coarse to ensure truly efficient spatial planning of no-go areas at smaller spatial scales, which would still require detailed, local-scale environmental impact assessments.

### 4.5 Conclusions

The rapid expansion of renewable energy facilities is essential to achieve global climate goals. However, this expansion needs careful management to avoid exacerbating the loss of biodiversity, particularly in regions that support threatened species. In many countries, such as Spain, PV infrastructure is being developed in highly suitable areas for steppe birds. Our analyses exemplify the challenges faced by wildlife adapted to open, flat, low productivity areas under a scenario of swift solar PV development, and how spatial prioritization may help reconcile climate change mitigation and biodiversity conservation goals. Our findings provide valuable insights for policymakers, energy developers, and conservationists, offering a framework for balancing renewable energy expansion with biodiversity conservation.

Also, our framework is applicable to other regions and species. For example, the use of comprehensive global trait databases (necessary for functional diversity analyses) available for different taxa such as birds (Tobias et al., 2022), mammals (Soria et al., 2021), amphibians (Oliveira et al., 2017), and reptiles (Oskyrko et al., 2024) can facilitate assessments of the impact of other infrastructures, such as roads (Rahhal et al., 2023) and wind farms (Thaxter et al., 2017). This is crucial in biodiversity hotspots, where thousands of species may be affected by human development (Medrano-Vizcaíno et al., 2022), yet distribution and trait data can be incomplete or incorrectly assumed for some species (e.g., Medrano-Vizcaíno & Rueda 2018).

By using an approach based on taxonomic, functional, and phylogenetic diversity to assess potential conflicts with human infrastructure, we can better understand the ecological consequences of human-induced environmental changes and develop strategies that promote both sustainability and biodiversity conservation.

## Data availability

The R code, and spatial data from TD, FD and PD analyses are available at: https://github.com/pabmevi/BiodiversityFacets_ConsBio. The trait data is provided as supplementary material. Occurrence data for each species can be made available upon request to the authors and SEO/ BirdLife.

## Acknowledgements

This study was funded by the ELECTROSTEPPE Project (TED2021-130352B-I00 funded by MCIN/AEI/10.13039/ 501100011033 and the EU “NextGenerationEU”/ PRTR). This work contributes to the Steppe-Forward Chair (UAM-CTFC-TotalEnergies) and the project 022-GRIN-34462 awarded by the University of Castilla-La Mancha & Fondo Europeo de Desarrollo Regional (FEDER). We are grateful to SEO/Birdlife for contributing data from the II and III Spanish Atlas of Breeding Birds. ABL acknowledges financial support by a Ramón y Cajal grant (RYC2021-031737-I) funded by MCIN/AEI/10.13039/ 501100011033 and the EU “NextGenerationEU”/PRTR. This research benefited from project 2021-SGR 00302 funded by the Departament de Recerca i Universitats de la Generalitat de Catalunya.

## Notes

https://github.com/pabmevi/BiodiversityFacets_ConsBio

### Competing Interest Statement

The authors have declared no competing interest.

## References

Aros-Mualín, D., Noben, S., Karger, D. N., Carvajal-Hernández, C. I., & Guisan, A. (2021). Functional diversity in ferns is driven by species richness rather than by environmental constraints. Frontiers in Plant Science, 11, 615723. 10.3389/fpls.2020.615723

Bastani, F., Wolters, P., Gupta, R., Ferdinando, J., & Kembhavi, A. (2023). SatlasPretrain: A large-scale dataset for remote sensing image understanding. Proceedings of the IEEE International Conference on Computer Vision, 2023, 16726–16736. 10.1109/ICCV51070.2023.01538

Boakes, E. H., Fuller, R. A., & McGowan, P. J. K. (2019). The extirpation of species outside protected areas. Conservation Letters, 12, e12608. 10.1111/conl.12608

BOE (2023). Real Decreto 445/2023, de 13 de junio, por el que se modifican los Anexos I, II y III de la Ley 21/2013, de 9 de diciembre, de evaluación ambiental. Boletín Oficial del Estado, 140, 87720–87735. https://www.boe.es/eli/es/rd/2023/06/13/445

Burfield, I. J. (2005). The conservation status of steppic birds in Europe. In G. Bota, M. Morales, S. Mañosa, & J. Camprodon (Eds.), Ecology and conservation of steppe-land birds (pp. 119–139). Lynx Edicions & Centre Tecnològic Forestal de Catalunya. Barcelona, Spain

Burfield, I. J., Rutherford, C. A., Fernando, E., Gaget, E., Kalas, M., Keller, V., Knaus, P., & Nagy, S. (2023). Birds in Europe 4: The fourth assessment of species of European conservation concern. Bird Conservation International, 33, e71. 10.1017/S0959270923000187

Cadotte, M. W., & Tucker, C. M. (2018). Difficult decisions: Strategies for conservation prioritization when taxonomic, phylogenetic and functional diversity are not spatially congruent. Biological Conservation, 225, 128–133. 10.1016/j.biocon.2018.06.014

Cadotte, M. W., Dinnage, R., & Tilman, D. (2012). Phylogenetic diversity promotes ecosystem stability. Ecology, 93, S223–S233. 10.1890/11-0426.1

CBD. (2010). COP 10 Decision X/2: Strategic plan for biodiversity 2011–2020. Convention on Biological Diversity. Montreal, Canada. https://www.cbd.int/decision/cop/?id=12268

Chien, F., Hsu, C. C., Ozturk, I., Sharif, A., & Sadiq, M. (2022). The role of renewable energy and urbanization towards greenhouse gas emission in top Asian countries: Evidence from advance panel estimations. Renewable Energy, 186, 207–216. 10.1016/j.renene.2021.12.118

Cooke, R. S., Eigenbrod, F., & Bates, A. E. (2019). Projected losses of global mammal and bird ecological strategies. Nature Communications, 10, 2279. 10.1038/s41467-019-10284-z

Copping, J. P., Waite, C. E., Balmford, A., Bradbury, R. B., Field, R. H., Morris, I., & Finch, T. (2025). Solar farm management influences breeding bird responses in an arable-dominated landscape. Bird Study, 72, 1–6. 10.1080/00063657.2025.2450392

Dhar, A., Naeth, M. A., Jennings, P. D., & El-Din, M. G. (2020). Perspectives on environmental impacts and a land reclamation strategy for solar and wind energy systems. Science of the Total Environment, 718, 134602. 10.1016/j.scitotenv.2019.134602

Dunnett, S., Holland, R. A., Taylor, G., & Eigenbrod, F. (2022). Predicted wind and solar energy expansion has minimal overlap with multiple conservation priorities across global regions. Proceedings of the National Academy of Sciences USA, 119, e2104764119. 10.1073/pnas.2104764119

European Commission. (2019). The European Green Deal: Communication from the Commission to the European Parliament, the European Council, the Council, the European Economic and Social Committee and the Committee of the Regions. European Commission. Brussels, Belgium

European Commission. (2020). EU Biodiversity Strategy for 2030. European Commission. Brussels, Belgium. https://environment.ec.europa.eu/strategy/biodiversity-strategy-2030_en

European Commission. (2022). REPowerEU Plan: Communication from the Commission to the European Parliament, the European Council, the Council, the European Economic and Social Committee and the Committee of the Regions. European Commission. Brussels, Belgium

European Parliament and Council of the European Union. (2018). Directive (EU) 2018/2001 of the European Parliament and of the Council of 11 December 2018 on the promotion of the use of energy from renewable sources. Official Journal of the European Union, L 328, 82–209. https://eur-lex.europa.eu/eli/dir/2018/2001/oj

Faith, D. P. (1992). Conservation evaluation and phylogenetic diversity. Biological Conservation, 61, 1–10. 10.1016/0006-3207(92)91201-3

Fischer, A., Sandström, C., Delibes-Mateos, M., Arroyo, B., Tadie, D., Randall, D., Hailu, F., Lowassa, A., Msuha, M., Kereži, V., Reljić, S., Linnell, J., & Majić, A. (2013). On the multifunctionality of hunting: An institutional analysis of eight cases from Europe and Africa. Journal of Environmental Planning and Management, 56, 531–552. 10.1080/09640568.2012.689615

Gómez-Catasús, J., Morales, M. B., Giralt, D., González del Portillo, D., Manzano-Rubio, R., Solé-Bujalance, L., Sardá-Palomera, F., Traba, J. & Bota, G., (2024). Solar photovoltaic energy development and biodiversity conservation: Current knowledge and research gaps. Conservation Letters, 17, e13025. 10.1111/conl.13025

Gómez-Catasús, J., Pérez-Granados, C., Barrero, A., Bota, G., Giralt, D., López-Iborra, G. M., Serrano, D., & Traba, J. (2018). European population trends and current conservation status of an endangered steppe-bird species: The Dupont’s lark Chersophilus duponti. PeerJ, 6, e5627. 10.7717/peerj.5627

Gómez-Catasús, J., Benítez-López, A., Díaz, M., et al (2025). Alarming conservation status of Western European steppe birds and their habitats: An expert-based review of current threats, traits and knowledge gaps. Biological Conservation, 311, 111414. 10.1016/j.biocon.2025.111414

Guerrero, I., Duque, D., Oñate, J. J., Pärt, T., Bengtsson, J., Tscharntke, T., Liira, J., Aavik, T., Emmerson, M., Berendse, F., Ceryngier, P., Weisser, W., & Morales, M. B. (2024). Agricultural intensification affects birds’ trait diversity across Europe. Basic and Applied Ecology, 74, 40–48. 10.1016/j.baae.2023.11.007

Guil, F., & Pérez-García, J. M. (2022). Bird electrocution on power lines: Spatial gaps and identification of driving factors at global scales. Journal of Environmental Management, 301, 113890. 10.1016/j.jenvman.2021.113890

Hermoso, V., Bota, G., Brotons, L., & Mor, A. (2023). Land use policy addressing the challenge of photovoltaic growth: Integrating multiple objectives towards sustainable green energy development. Land Use Policy, 128, 106592. 10.1016/j.landusepol.2023.106592

IEA. (2021). Energy Policy Review Spain 2021. International Energy Agency. Paris, France. https://iea.blob.core.windows.net/assets/2f405ae0-4617-4e16-884c-7956d1945f64/Spain2021.pdf

Instituto Geográfico Nacional. (n.d.). Plan Nacional de Ortofotografía Aérea (PNOA). Retrieved 25 Sept. 2024, https://pnoa.ign.es/web/portal/inicio

IPBES. (2019). Global assessment report on biodiversity and ecosystem services. IPBES Secretariat. Bonn, Germany

Jetz, W., Thomas, G., Joy, J., Hartmann, K., & Mooers, A. O. (2012). The global diversity of birds in space and time. Nature, 491, 444–448. 10.1038/nature11631

Johnson, T. F., Isaac, N. J. B., Paviolo, A., & González-Suárez, M. (2021). Handling missing values in trait data. Global Ecology and Biogeography, 30, 51–62. 10.1111/geb.13185

Kembel, S., Cowan, P., Helmus, M., Cornwell, H., Morlon, H., Ackerly, D., Blomberg, S., & Webb, C. (2010). Picante: R tools for integrating phylogenies and ecology. Bioinformatics, 26, 1463–1464. 10.1093/bioinformatics/btq166

Kruitwagen, L., Story, K. T., Friedrich, J., Byers, L., Skillman, S., & Hepburn, C. (2021). A global inventory of photovoltaic solar energy generating units. Nature, 598, 604–610. 10.1038/s41586-021-03957-7

Kunming-Montreal Global Biodiversity Framework. (2022). Kunming-Montreal Global Biodiversity Framework. Convention on Biological Diversity. Montreal, Canada. https://www.cbd.int/gbf

Lafitte, A., Sordello, R., Ouédraogo, D. Y., Thierry, C., Marx, G., Froidevaux, J., Schatz, B., Kerbiriou, C., Gourdain, P., & Reyjol, Y. (2023). Existing evidence on the effects of photovoltaic panels on biodiversity: A systematic map with critical appraisal of study validity. Environmental Evidence, 12, 25. 10.1186/s13750-023-00318-x

Laliberté, E., Legendre, P., & Shipley, B. (2014). FD: Measuring functional diversity from multiple traits, and other tools for functional ecology. R package version 1.0–12

Lambert, Q., Gros, R., & Bischoff, A. (2022). Ecological restoration of solar park plant communities and the effect of solar panels. Ecological Engineering, 182, 106722. 10.1016/j.ecoleng.2022.106722

La Sorte, F. A., Lepczyk, C. A., Aronson, M. F. J., Goddard, M. A., Hedblom, M., Katti, M., MacGregor-Fors, I., Mörtberg, U., Nilon, C. H., Warren, P. S., Williams, N. S. G., & Yang, J. (2018). The phylogenetic and functional diversity of regional breeding bird assemblages is reduced and constricted through urbanization. Diversity and Distributions, 24, 928–938. 10.1111/ddi.12738

Liu, Y., Zhang, R. Q., Huang, Z., Cheng, Z., López-Vicente, M., Ma, X. R., & Wu, G. L. (2019). Solar photovoltaic panels significantly promote vegetation recovery by modifying the soil surface microhabitats in an arid sandy ecosystem. Land Degradation and Development, 30, 2177–2186. 10.1002/ldr.3408

Lum, D., Rheindt, F. E., & Chisholm, R. A. (2022). Tracking scientific discovery of avian phylogenetic diversity over 250 years. Proceedings of the Royal Society B, 289, 20220088. 10.1098/rspb.2022.0088

Madroño, A., González, C., & Atienza, J. C. (2004). Libro rojo de las aves de España. Dirección General para la Biodiversidad-SEO/BirdLife. Madrid, Spain

Maire, E., Grenouillet, G., Brosse, S., & Villéger, S. (2015). How many dimensions are needed to accurately assess functional diversity? A pragmatic approach for assessing the quality of functional spaces. Global Ecology and Biogeography, 24, 728–740. 10.1111/geb.12299

Mason, N. W. H., De Bello, F., Mouillot, D., Pavoine, S., & Dray, S. (2013). A guide for using functional diversity indices to reveal changes in assembly processes along ecological gradients. Journal of Vegetation Science, 24, 794–806. 10.1111/jvs.12013

Mazel, F., Pennell, M. W., Cadotte, M. W., Diaz, S., Dalla Riva, G. V., Grenyer, R., Leprieur, F., Mooers, A. O., Mouillot, D., Tucker, C. M., & Pearse, W. D. (2018). Prioritizing phylogenetic diversity captures functional diversity unreliably. Nature Communications, 9, 2888. 10.1038/s41467-018-05126-3

Medrano-Vizcaíno, P., Grilo, C., Brito-Zapata, D., & González-Suárez, M. (2023). Landscape and road features linked to wildlife mortality in the Amazon. Biodiversity and Conservation, 32, 4337–4352. 10.1007/s10531-023-02699-4

Medrano-Vizcaíno, P., Benítez-López, A., Traba, J., Arroyo, B., Bota, G., Morales, M. B., & Mougeot, F. (2025). Spatial shifts in steppe bird hotspots over two decades: Assessing conservation priorities and the role of protected areas. Biological Conservation, 305, 111068. 10.1016/j.biocon.2025.111068

Medrano-Vizcaíno, P., Grilo, C., Silva Pinto, F. A., Carvalho, W. D., Melinski, R. D., Schultz, E. D., & González-Suárez, M. (2022). Roadkill patterns in Latin American birds and mammals. Global Ecology and Biogeography, 31(9), 1756–1783.

Medrano-Vizcaíno, P., & Rueda, A. (2018). Nuevo registro altitudinal del Pavón Nocturno *Nothocrax urumutum* (Cracidae) y notas sobre su historia natural. Revista Ecuatoriana de Ornitología, 3, 15–19.

MITECO (n.d.). Informes de instalaciones de producción de energía eléctrica. Accessed 25 Sept. 2024, https://energia.serviciosmin.gob.es/Pretor/Vista/Informes/InformesInstalaciones.aspx

MITECO (2024). Plan Nacional Integrado de Energía y Clima: Actualización 2023– 2030. Ministerio para la Transición Ecológica y el Reto Demográfico. Madrid, Spain. https://www.miteco.gob.es/es/prensa/pniec.aspx

Moilanen, A., Lehtinen, P., Kohonen, I., Jalkanen, J., Virtanen, E. A., & Kujala, H. (2022). Novel methods for spatial prioritization with applications in conservation, land use planning and ecological impact avoidance. Methods in Ecology and Evolution, 13, 1062–1072. 10.1111/2041-210X.13819

Monnet, A. C., Jiguet, F., Meynard, C. N., Mouillot, D., Mouquet, N., Thuiller, W., & Devictor, V. (2014). Asynchrony of taxonomic, functional and phylogenetic diversity in birds. Global Ecology and Biogeography, 23, 780–788. 10.1111/geb.12179

Moscatelli, M. C., Marabottini, R., Massaccesi, L., & Marinari, S. (2022). Soil properties changes after seven years of ground mounted photovoltaic panels in Central Italy coastal area. Geoderma Regional, 29, e00500. 10.1016/j.geodrs.2022.e00500

Niebuhr, B. B., & Sant, D. (2022). Renewable energy infrastructure impacts biodiversity beyond the area it occupies. Proceedings of the National Academy of Sciences USA, 119, e2200539119. 10.1073/pnas.2208815119

Oliveira, B. F., São-Pedro, V. A., Santos-Barrera, G., Penone, C., & Costa, G. C. (2017). AmphiBIO, a global database for amphibian ecological traits. Scientific data, 4(1), 1–7. 10.1038/sdata.2017.123

Oskyrko, O., Mi, C., Meiri, S., & Du, W. (2024). ReptTraits: a comprehensive dataset of ecological traits in reptiles. Scientific Data, 11(1), 243. 10.1038/s41597-024-03079-5

Penjor, U., Cushman, S. A., Kaszta, Ż. M., Sherub, S., & Macdonald, D. W. (2022). Effects of land use and climate change on functional and phylogenetic diversity of terrestrial vertebrates in a Himalayan biodiversity hotspot. Diversity and Distributions, 28, 2931–2943. 10.1111/ddi.13613

Pérez-García, J. M., Morant, J., Arrondo, E., Sebastián-González, E., Lambertucci, S. A., Santangeli, A., Margalida, A., Sánchez-Zapata, J., Donázar, J. A., Blanco, G., Donasar, J., Carrete, M. & Serrano, D. (2022). Priority areas for conservation alone are not a good proxy for predicting the impact of renewable energy expansion. Proceedings of the National Academy of Sciences USA, 119, e2204505119. 10.1073/pnas.2204505119

Pérez-Granados, C., López-Iborra, G. M., Serrano-Davies, E., Noguerales, V., Garza, V., Justribó, J. H., & Suárez, F. (2013). Short-term effects of a wildfire on the endangered Dupont’s lark Chersophilus duponti in an arid shrub-steppe of central Spain. Acta Ornithologica, 48, 201–210. 10.3161/000164513X678856

Phalan, B., Hayes, G., Brooks, S., Marsh, D., Howard, P., Costelloe, B., … & Whitaker, S. (2018). Avoiding impacts on biodiversity through strengthening the first stage of the mitigation hierarchy. Oryx, 52(2), 316–324. 10.1017/S0030605316001034

Plass-Johnson, J. G., Taylor, M. H., Husain, A., Teichberg, M. & Ferse, S. (2016). Non-random variability in functional composition of coral reef fish communities along an environmental gradient. PLoS ONE, 11, e0154014. 10.1371/journal.pone.0154014

Pollock, L. J., Thuiller, W., & Jetz, W. (2017). Large conservation gains possible for global biodiversity facets. Nature, 546, 141–144. 10.1038/nature22368

Pourasl, H. H., Vatankhah, R., & Khojastehnezhad, V. M. (2023). Solar energy status in the world: A comprehensive review. Energy Reports, 10, 3474–3493. 10.1016/j.egyr.2023.10.022

Powers, R. P., & Jetz, W. (2019). Global habitat loss and extinction risk of terrestrial vertebrates under future land-use-change scenarios. Nature Climate Change, 9, 323–329. 10.1038/s41558-019-0406-z

Prener, C., Grossenbacher, T., & Zehr, A. (2022). biscale: Tools and palettes for bivariate thematic mapping. R package version 1.0.0. https://CRAN.R-project.org/package=biscale

Red Eléctrica de España. (2025). Potencia instalada. Red Eléctrica de España. Madrid, Spain. https://www.sistemaelectrico-ree.es/informe-del-sistema-electrico/potencia-instalada

Romero-Muñoz, A., Fandos, G., Benítez-López, A., & Kuemmerle, T. (2021). Habitat destruction and overexploitation drive widespread declines in all facets of mammalian diversity in the Gran Chaco. Global Change Biology, 27, 755–767. 10.1111/gcb.15418

Rosauer, D., Laffan, S. W., Crisp, M. D., Donnellan, S. C., & Cook, L. G. (2009). Phylogenetic endemism: A new approach for identifying geographical concentrations of evolutionary history. Molecular Ecology, 18, 4061–4072. 10.1111/j.1365-294X.2009.04311.x

Sainz Ollero, H., van Staalduinen, M.A., 2012. Iberian steppes. In: Eurasian Steppes: Ecological Problems and Livelihoods in a Changing World, pp. 273–288. 10.1007/978-94-007-3886-7_9.

Santangeli, A., Minin, E., Toivonen, T., & Pogson, M. (2016). Synergies and trade-offs between renewable energy expansion and biodiversity conservation: A cross-national multifactor analysis. GCB Bioenergy, 8, 1191–1200. 10.1111/gcbb.12337

Santini, L., Tobias, J. A., Callaghan, C., Gallego-Zamorano, J., & Benítez-López, A. (2023). Global patterns and predictors of avian population density. Global Ecology and Biogeography, 32, 1189–1204. 10.1111/geb.13688

Santos, T., & Suárez, F. (2005). Biogeography and population trends of Iberian steppe birds. In G. Bota, M. B. Morales, S. Mañosa, & J. Camprodon (Eds.), Ecology and conservation of steppe-land birds (pp. 69–102). Lynx Edicions & Centre Tecnològic Forestal de Catalunya. Barcelona, Spain

Sanz-Pérez, A., Giralt, D., Robleño, I., Bota, G., Milleret, C., Mañosa, S., & Sardà-Palomera, F. (2019). Fallow management increases habitat suitability for endangered steppe bird species through changes in vegetation structure. Journal of Applied Ecology, 56, 2166–2175. 10.1111/1365-2664.13450

Semeraro, T., Scarano, A., Santino, A., Rohinton, E. & Lenucci, M. (2022). An innovative approach to combine solar photovoltaic gardens with agricultural production and ecosystem services. Ecosystem Services, 56, 101450. 10.1016/j.ecoser.2022.101450

SEO/BirdLife. (2021). Libro rojo de las aves de España. SEO/BirdLife. Madrid, Spain

SEO/BirdLife, Molina, B., Nebreda, A., Muñoz, A. R., Seoane, J., Real, R., Bustamante, J., & Del Moral, J. C. (2022). III Atlas de aves en época de reproducción en España. SEO/BirdLife. Madrid, Spain. https://atlasaves.seo.org

Serrano, D., Margalida, A., Pérez-García, J. M., Juste, J., Traba, J., Valera, F., Carrete, M., Aihartza, J., Real, J., ….. & Mañosa, S. (2020). Renewables in Spain threaten biodiversity. Science, 370, 1282–1283. 10.1126/science.abf6509

Smiley, T. M., Title, P. O., Zelditch, M. L., & Terry, R. C. (2020). Multi-dimensional biodiversity hotspots and the future of taxonomic, ecological and phylogenetic diversity: A case study of North American rodents. Global Ecology and Biogeography, 29, 516–533. 10.1111/geb.13050

Snow, D. W., & Perrins, C. M. (Eds.). (1998). The birds of the western Palearctic: Concise edition. Oxford University Press. Oxford, UK

Sol, D., Trisos, C., Múrria, C., Jeliazkov, A., González-Lagos, C., Pigot, A., Ricotta, C., Swan, C., Tobias, J. & Pavoine, S. (2020). The worldwide impact of urbanisation on avian functional diversity. Ecology Letters, 23, 962–972. 10.1111/ele.13495

Sonter, L. J., Dade, M. C., Watson, J. E. M., & Valenta, R. K. (2020). Renewable energy production will exacerbate mining threats to biodiversity. Nature Communications, 11, 4174. 10.1038/s41467-020-17928-5

Stekhoven, D. (2013). missForest: Nonparametric missing value imputation using Random Forest. R package version 1.4

Sturchio, M. A., & Knapp, A. K. (2023). Ecovoltaic principles for a more sustainable, ecologically informed solar energy future. Nature Ecology and Evolution, 7, 1746–1749. 10.1038/s41559-023-02174-x

Suárez, F., Naveso, M. A., & De Juana, E. (1997). Farming in the drylands of Spain: Birds of the pseudosteppes. In D. J. Pain & N. W. Pienkowski (Eds.), Farming and birds in Europe: The common agricultural policy and its implications for bird conservation (pp. 79–116). Academic Press. San Diego, CA

Thaxter, C. B., Buchanan, G. M., Carr, J., Butchart, S. H., Newbold, T., Green, R. E., … & Pearce-Higgins, J. W. (2017). Bird and bat species’ global vulnerability to collision mortality at wind farms revealed through a trait-based assessment. Proceedings of the Royal Society B: Biological Sciences, 284(1862), 20170829. 10.1098/rspb.2017.0829

Tobias, J. A., Sheard, C., Pigot, A. L., Devenish, A. J. M., Yang, J., Sayol, F., Neate-Clegg., ..… & Schleuning, M. (2022). AVONET: Morphological, ecological and geographical data for all birds. Ecology Letters, 25, 581–597. 10.1111/ele.13898

Tölgyesi, C., Bátori, Z., Pascarella, J., Erdős, L., Török, P. & Gallé, R (2023). Ecovoltaics: Framework and future research directions to reconcile land-based solar power development with ecosystem conservation. Biological Conservation, 285, 110242. 10.1016/j.biocon.2023.110242

Traba, J., & Morales, M. B. (2019). The decline of farmland birds in Spain is strongly associated to the loss of fallowland. Scientific Reports, 9, 9473. 10.1038/s41598-019-45854-0

Traba, J., & Pérez-Granados, C. (2022). Extensive sheep grazing is associated with trends in steppe birds in Spain: Recommendations for the Common Agricultural Policy. PeerJ, 10, e12870. 10.7717/peerj.12870

Traba, J., García de la Morena, E. L., Morales, M. B., & Suárez, F. (2007). Determining high value areas for steppe birds in Spain: Hotspots, complementarity and the efficiency of protected areas. Biodiversity and Conservation, 16, 3255–3275. 10.1007/s10531-006-9138-2

Tsuboi, M., van der Bijl, W., Kopperud, B. T., Erritzoe, J., Voje, K. L., Kotrschal, A., Yopak, K. E., Collin, S. P., Iwaniuk, A. N., & Kolm, N. (2018). Breakdown of brain–body allometry and the encephalization of birds and mammals. Nature Ecology and Evolution, 2, 1492–1500. 10.1038/s41559-018-0632-1

Tucker, C. M., Cadotte, M. W., Carvalho, S. B., Davies, T. J., Ferrier, S., Fritz, S. A., Grenyer, R., Helmus., ….. & Mazel, F. (2017). A guide to phylogenetic metrics for conservation, community ecology and macroecology. Biological Reviews, 92, 698–715. 10.1111/brv.12252

van de Ven, D. J., Capellan-Peréz, I., Arto, I., Cazcarro, I., de Castro, C., Patel, P. & González-Eguino, M. (2021). The potential land requirements and related land use change emissions of solar energy. Scientific Reports, 11, 2907. 10.1038/s41598-021-82042-5

Vermeylen, S. (2010). Resource rights and the evolution of renewable energy technologies. Renewable Energy, 35, 2399–2405. 10.1016/j.renene.2010.03.017

Villéger, S., Mason, N. W. H., & Mouillot, D. (2008). New multidimensional functional diversity indices for a multifaceted framework in functional ecology. Ecology, 89, 2290–2301. 10.1890/07-1206.1

Visser, E., Perold, V., Ralston-Paton, S., Cardenal, A. C., & Ryan, P. G. (2019). Assessing the impacts of a utility-scale photovoltaic solar energy facility on birds in the Northern Cape, South Africa. Renewable Energy, 133, 1285–1294. 10.1016/j.renene.2018.08.106

Walston, L. J., Rollins, K. E., LaGory, K. E., Smith, K. P., & Meyers, S. A. (2016). A preliminary assessment of avian mortality at utility-scale solar energy facilities in the United States. Renewable Energy, 92, 405–414. 10.1016/j.renene.2016.02.041

Wilson, K. A., & Law, E. A. (2016). How to avoid underselling biodiversity with ecosystem services: A response to Silvertown. Trends in Ecology and Evolution, 31, 332–333. 10.1016/j.tree.2016.03.002

