## Supplementary material for "Identifying the exposure of taxonomic, functional, and phylogenetic diversity of steppe birds to renewable energy developments"

**Table S1.** Eigenvalues and their contribution to the functional space quality used to select the number of PCoA dimensions

| **Eigenvalues** | **Relative_eig** | **Rel_corr_eig** | **Broken_stick** | **Cum_corr_eig** | **Cumul_br_stick** |
| --- | --- | --- | --- | --- | --- |
| 0.798 | 0.532 | 0.371 | 0.157 | 0.371 | 0.157 |
| 0.268 | 0.179 | 0.133 | 0.116 | 0.504 | 0.273 |
| 0.146 | 0.098 | 0.079 | 0.095 | 0.583 | 0.368 |
| 0.103 | 0.069 | 0.059 | 0.081 | 0.642 | 0.449 |
| 0.093 | 0.062 | 0.055 | 0.071 | 0.696 | 0.519 |
| 0.065 | 0.043 | 0.042 | 0.062 | 0.739 | 0.581 |
| 0.039 | 0.026 | 0.030 | 0.055 | 0.769 | 0.637 |
| 0.037 | 0.025 | 0.030 | 0.049 | 0.799 | 0.686 |
| 0.024 | 0.016 | 0.024 | 0.044 | 0.823 | 0.730 |
| 0.018 | 0.012 | 0.021 | 0.039 | 0.84 | 0.770 |

**Table S2**. Pearson correlation matrix of TD, FD, SESFD, PD, and SESPD.

| **Indexes** | **TD** | **FRic** | **SESFRic** | **PD** | **SESPD** |
| --- | --- | --- | --- | --- | --- |
| **TD** | 1.0 | 0.8 | 0.3 | 0.9 | -0.1 |
| **FRic** | 0.8 | 1.0 | 0.6 | 0.8 | 0.2 |
| **SESFRic** | 0.3 | 0.6 | 1.0 | 0.4 | 0.5 |
| **PD** | 0.9 | 0.8 | 0.4 | 1.0 | 0.3 |
| **SESPD** | -0.1 | 0.2 | 0.5 | 0.3 | 1.0 |

**Table S3.** Threat status categories used for analyses, definition for each category, and numerical scores assigned according to threat categories.

| **Criteria** | **Category** | **Category definition** | **Score assigned** |
| --- | --- | --- | --- |
| European population status (EPS) 2023 | EN | European population meets any of the IUCN Red List criteria for EN (Burfield et al. 2023). | 10 |
|  | VU | European population meets any of the IUCN Red List criteria for VU (Burfield et al. 2023). | 7 |
|  | NT | European population approaches the IUCN Red List criteria for VU (Burfield et al. 2023). | 6 |
|  | Declining | European population has declined by ≥20% since c.1980 and has continued to decline since c.2007; or trend since c.1980 unknown or uncertain, but European population has declined by ≥20% since c.2007; or European range contracted between the atlases (i.e. range change index value ≤-5) and European population has continued to decline since c.2007 (Burfield et al. 2023). | 5 |
|  | Rare | European population is <10,000 breeding pairs (or <30,000 wintering individuals) and is not marginal to a larger non-European population (Burfield et al. 2023). | 3 |
|  | Depleted | European population has declined by ≥20% since c.1980, but is not known or thought to have declined further since c.2007; or European range contracted between the atlases (i.e. range change index value ≤-5), but European population is not known or thought to have declined further since c.2007 (Burfield et al. 2023). | 2 |
|  | SecureF | European population does not meet any of the criteria above, but formerly qualified as a SPEC in one or more previous assessments and may not yet have fully recovered to its former population level or range extent (Burfield et al. 2023). | 1 |
|  | Secure | European population does not meet any of the criteria above (Burfield et al. 2023). | 0 |
| Species of European Conservation Concern (SPEC) 2023 | 1 | Species of global conservation concern, i.e. classified as Critically Endangered, Endangered, Vulnerable or Near Threatened at global level (BirdLife International 2022). | 10 |
|  | 2 | Species whose global population is concentrated in Europe, and which is classified as Regionally Extinct, Critically Endangered, Endangered, Vulnerable or Near Threatened at European level (BirdLife International 2021), or as Declining, Depleted or Rare in Europe (Burfield et al. 2023). | 7 |
|  | 3 | Species whose global population is not concentrated in Europe, but which is classified as Regionally Extinct, Critically Endangered, Endangered, Vulnerable or Near Threatened at European level (BirdLife International 2021) (unless it is marginal in Europe, not decreasing and qualifies solely under Criterion D; IUCN 2012a), or as Declining, Depleted or Rare in Europe (Burfield et al. 2023). | 5 |
|  | Non-SPEC | Species whose global population is not concentrated in Europe, and whose European population status is currently considered to be Secure or SecureF (Burfield et al. 2023). | 1 |
|  | Non-SPECe | Species whose global population is concentrated in Europe, but whose European population status is currently considered to be Secure or SecureF (Burfield et al. 2023). | 0 |
| Spanish Catalogue of Protected and Threatened Species (SCPTS) 2023 | Endangered | A species, subspecies, or population of a species whose survival is unlikely if the causal factors of its current situation continue to operate. | 10 |
|  | Vulnerable | A species, subspecies, or population of a species that is at risk of moving to the previous category in the near future if the adverse factors affecting it are not corrected. | 6.7 |
|  | LESRPE | Species, subspecies, and populations deserving of particular attention and protection based on their scientific, ecological, cultural value, uniqueness, rarity, or degree of threat. | 3.3 |
|  | NO | Not included in this list | 0 |

**Table S4.** Retention percentage of biodiversity facets and threatened species areas under each prioritization scenario.

| ***Prioritization scenarios*** | **TD%** | **FD%** | **PD%** | **Threatened%** | **Average%** |
| --- | --- | --- | --- | --- | --- |
| ***TD*** | 100 | 50.1 | 33.9 | 73.3 | 64.3 |
| ***FD*** | 50.1 | 100 | 54.2 | 49.4 | 63.4 |
| ***PD*** | 33.9 | 54.2 | 100 | 37.1 | 56.3 |
| ***TD+FD*** | 77.9 | 72.2 | 44.8 | 70.84 | 66.4 |
| ***TD+PD*** | 74.3 | 58.9 | 59.6 | 69.47 | 65.6 |
| ***FD+PD*** | 43.2 | 81.5 | 72.7 | 45.18 | 60.6 |
| ***TD+FD+PD*** | 68.5 | 75.5 | 59.7 | 62.8 | 66.6 |

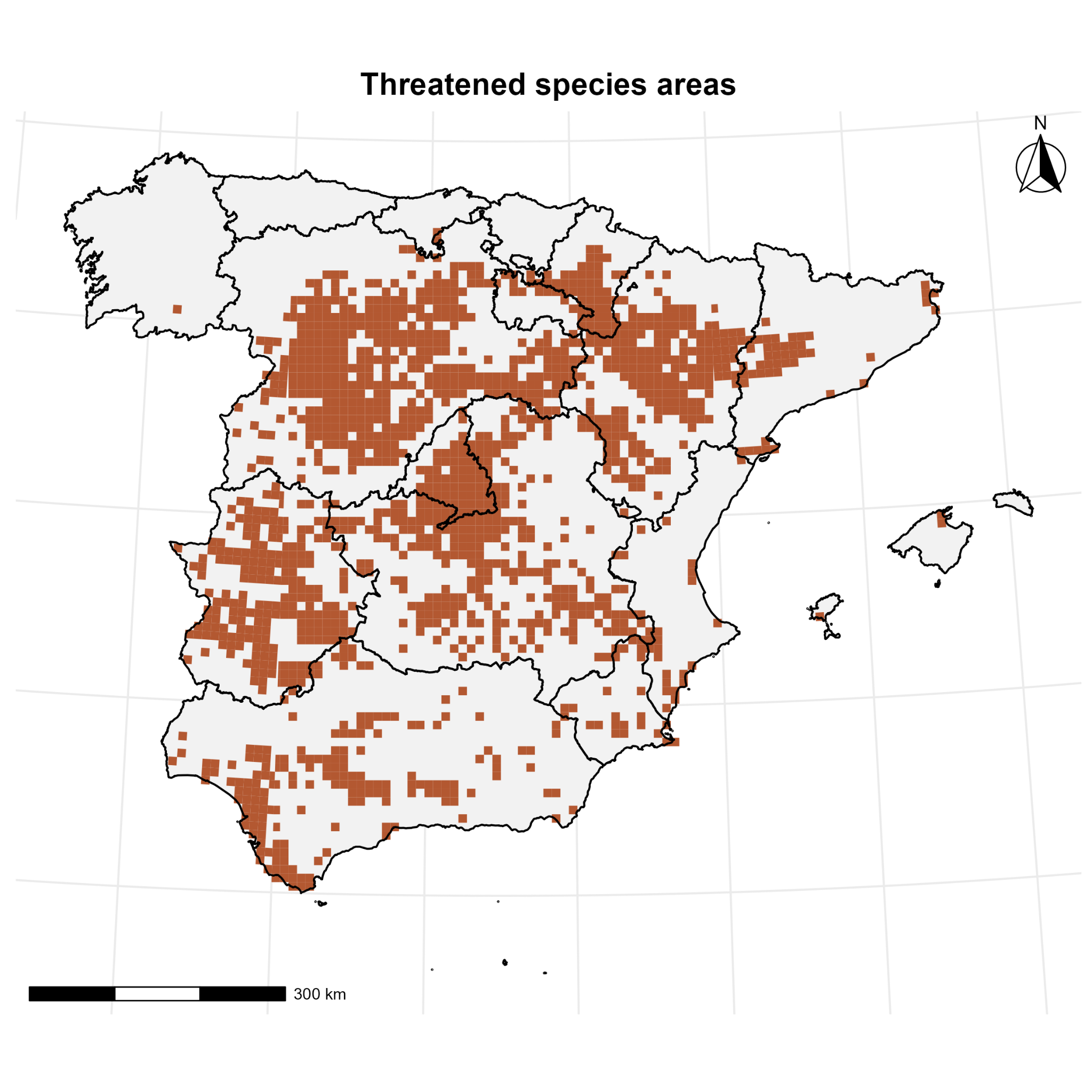

**Figure S1.** Threatened species index map based on the intersection of threat categories from the European Population Status (EPS), the Species of European Conservation Concern (SPEC), and the Spanish Catalogue of Threatened Species category (SCTS).

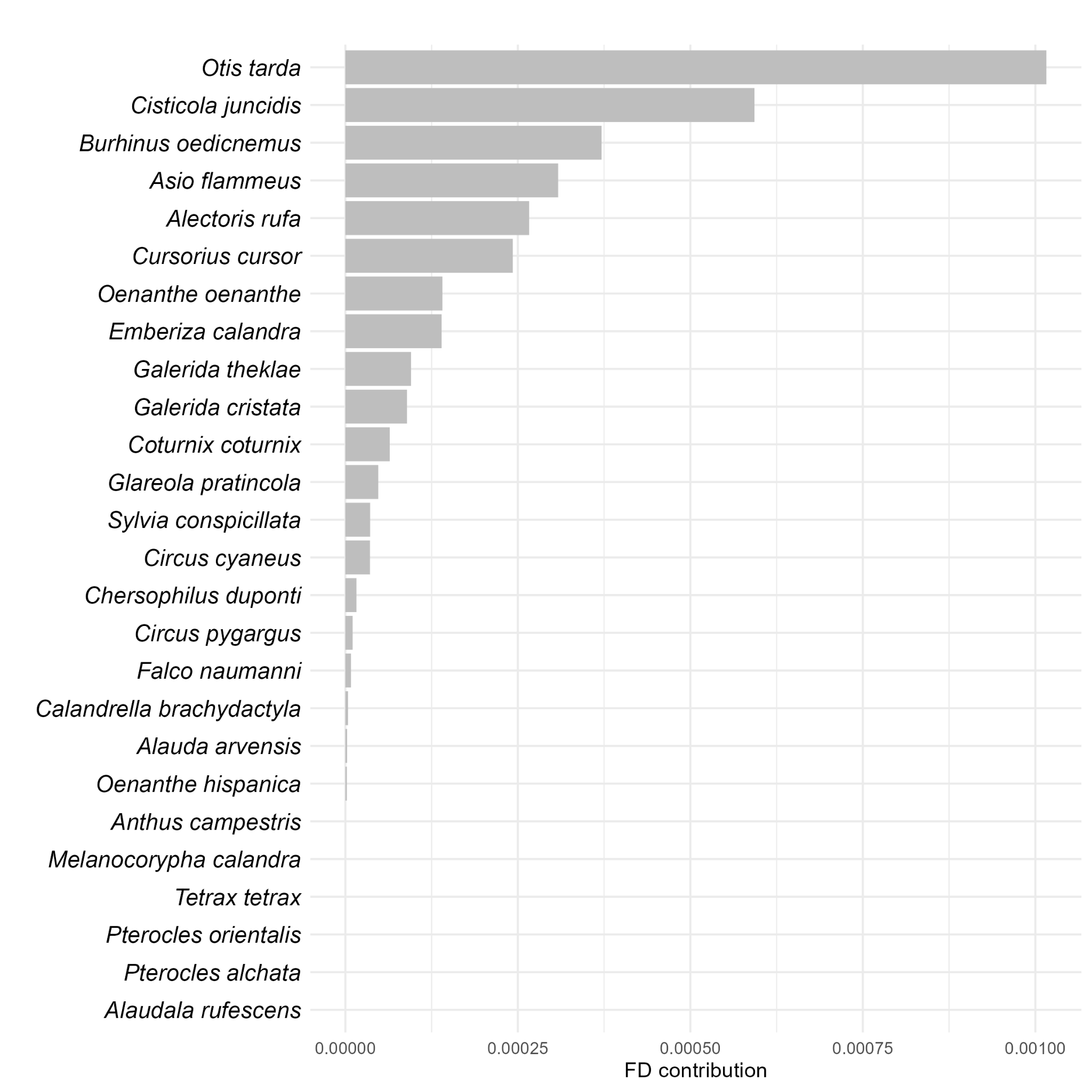

**Figure S2.** Species-level contribution to Functional Diversity, measured by the reduction in functional volume upon its individual exclusion.

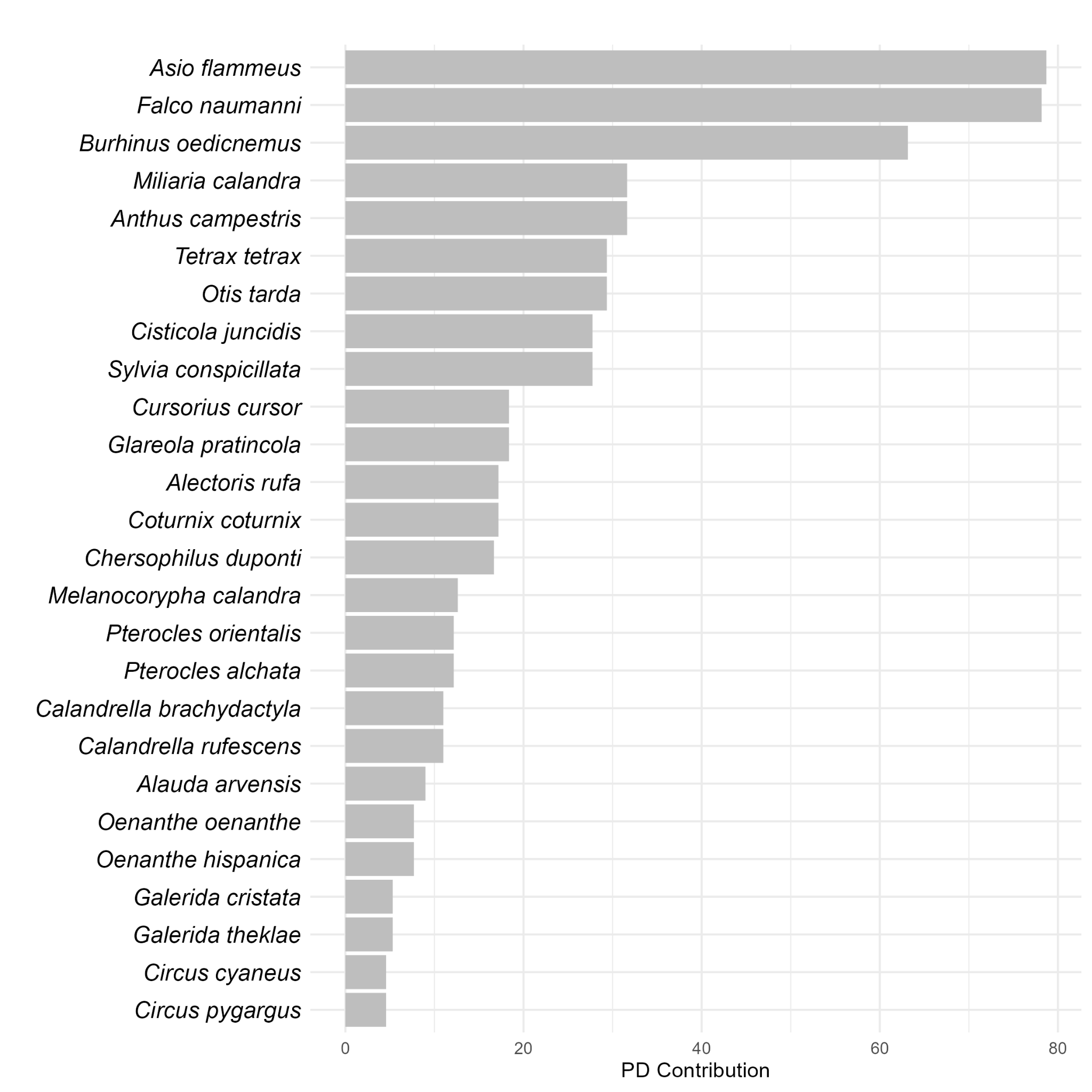

**Figure S3**. Species-level contribution to Phylogenetic Diversity, measured by the reduction in total branch length upon individual species exclusion.

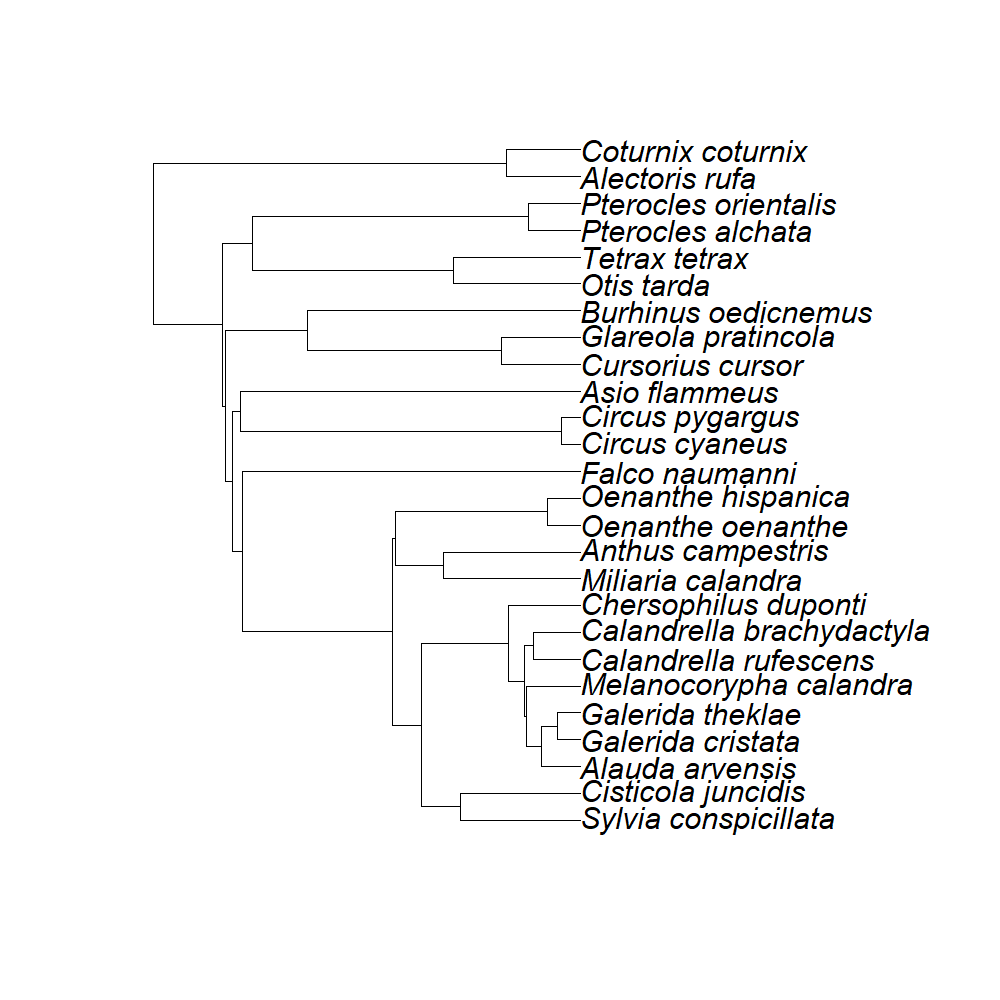

**Figure S4**. Phylogenetic relationships among the studied steppe bird species
